# G-quadruplex profiling in complex tissues using single-cell CUT&Tag

**DOI:** 10.1101/2025.04.03.647101

**Authors:** Jing Lyu, Szabolcs Hetey, Goncalo Castelo-Branco, Marek Bartosovic, Simon J Elsässer

## Abstract

G-quadruplexes (G4) are non-canonical DNA structures that gained increasing attention for their potential roles in gene regulation, with implications in neurodegenerative diseases and cancer. Despite their biological significance, G4 structures have not been studied systematically across tissues and cell types. In this study, we employ G4 single-cell CUT&Tag (G4 scCUT&Tag) to characterize G4 landscapes in postnatal mouse brain cells, leveraging single-cell analytical approaches commonly used in scRNA-Seq and scATAC-Seq datasets. Using conventional single-cell omics workflows to process and explore our data, we distinguished different cell lineages based on G4 heterogeneity and established that a subset of lineage-specific genes show unique promoter G4s. Multi-omics integration with scRNA-Seq gene expression profiles, using both a covariance-based technique (canonical correlation analysis) and a transfer learning-based approach, enabled a more detailed annotation of cell types. These integrations not only revealed significant correlation of G4 and gene expression signals, but demonstrated that G4 scCUT&Tag enables detailed examination of G4 heterogeneity in complex tissues and supports integrative analysis of G4 profiles with other omics layers, offering new insights into the epigenomic landscapes of the developing brain.

## Introduction

G-quadruplexes (G4) (1) (2) are the most extensively (3) (4) (5) investigated non-canonical conformations in living cells (6) (7), sparking interest in anticancer drug development (8) (9) (10) and understanding the molecular mechanisms of aging and age-related neurodegenerative disorders (11) (12). G4 landscapes differ substantially between cell types and can have prominent roles in cell type-specific processes (13), especially in the central nervous system (CNS) (14) (15) (16) (17). Therefore, the investigation of endogenous G4 landscapes in primary cells and tissues is a prerequisite for understanding the regulatory potential and disease impact of G4 structures. Recent single-cell methods revolutionize biology by enabling the measurement of cellular heterogeneity and discerning cell identity at unprecedented scale and resolution within cellular populations. As G4s are fundamental features of the chromatin and implicated in numerous important cellular processes such as transcription (18) (19) (20) (21), translation (22) (23) (24) and maintenance of genome stability (25) (26) (27), their single-cell profiling could lead to clearer understanding of the role G4s play in these mechanisms. The CUT&Tag methodology has been successfully used to map G4 structures in cells, including single cells (G4 scCUT&Tag) (28), but G4 landscapes of primary tissues have not been explored with this methodology. Here we perform G4 scCUT&Tag on the developing mouse brain and apply single-cell computational approaches like marker analysis, label imputation, linear-embedding (canonical correlation analysis (29,30)), deep learning-driven multi-omics integration (scBridge (31)) and cis-regulatory interaction predictions (cicero (32)). We find that G4 structures reveal cell-specific G4 signatures, and cell type-specific clustering can be greatly enhanced by integration with other modalities such as single-cell RNA-Seq (scRNA-Seq). Moreover, G4 scCUT&Tag data allows us to predict G4 regions of the genome that are more likely to be in physical proximity with each other in the nucleus.

## Results

### scCUT&Tag mapping of G-quadruplexes allows for distinguishing specific cell populations

In order to map G4s at the single-cell level we developed a highly sensitive centrifugation-based variant of our bulk G4 CUT&Tag (33) with a previously established scCUT&Tag protocol based on the droplet-based 10x Genomics scATAC kit (34) (**Fig 1A**). Bulk G4 CUT&Tag derived from centrifugation-based protocol showed superior signal-to-noise ratio and sensitivity to weaker G4 signals (**Suppl. Fig 1A-D**). To validate the single-cell G4 CUT&Tag workflow, we prepared a heterogeneous cell mix and attempted to resolve single cells into discrete groups recapitulating the original cell types based exclusively on G4 patterns. We mixed mouse embryonic stem cells (mESC, C57Bl/6J origin) with mouse embryonic fibroblasts (MEF, NIH-3T3) (ATCC), and following BG4 antibody-based in situ labeling and tagmentation, we performed scCUT&Tag using the 10x Genomics scATAC-Seq kit. After processing the single-cell data we obtained 1842 single cell profiles with 3434 median reads passing quality filter. The median fraction of reads in peaks (FRIP) was 52%, indicating a high peak enrichment characteristic of CUT&Tag data (33,35). In order to determine whether standard clustering approaches are able to identify cell types based on the G4 signal, we performed dimensional reduction by latent semantic indexing (LSI) and uniform manifold approximation and projection (UMAP) followed by shared nearest neighbors (SNN) driven clustering by Seurat package. Using this approach with resolution parameter 0.1 we obtained two distinguishable clusters (**Fig 1B**) with similar quality profiles (**Fig 1C**). To annotate these clusters, at first, we made an overlap analysis applied on MACS2 peak sets coming from bulk CUT&Tag experiments and validated that cluster 1 (mESC cluster hereafter) shows higher overlap (6,543/34,152 peaks) with high-confidence G4 sites of mESC G4 than cluster 0 (1,259/31,019 peaks). Consistent with this finding, cluster 0 (MEF cluster hereafter) had higher overlap (3,321/31,019 peaks) with the MEF G4 peak set than cluster 1 (1,113/34,152 peaks) (**Fig 1D**). As expected from prior observations that G4s mark many housekeeping gene promoters, we observed considerable overlap between MEF and mESC cluster peaks. In bulk mESC and MEF comparisons, 8,634 and 3,919 peaks were detected in both Seurat clusters, respectively. Additionally, a substantial number of peaks (12,422 out of 34,152 for mESC and 17,137 out of 31,019 for MEF) were found across all three sets. Principal component analysis (PCA) of the marker peaks showed strong co-clustering between the respective bulk and scCUT&Tag data (**Fig 1E**). Also, normalized G4 and mESC ATAC-Seq signals over the cluster-specific peaks from the overlap analysis confirmed the cluster annotations (**Fig 1F**). We further validated the Seurat clusters through genome-wide correlation analysis using our centrifugation-based bulk G4 CUT&Tag datasets. Pearson correlation coefficients demonstrated a clear similarity between the single-cell clusters and their bulk counterparts (**Fig 1G**). Lastly, representative MEF and mESC-specific G4 peaks were visualized by normalized genome browser tracks (**Fig 1H**) and coverage plots (**Suppl. Fig 1F**). These results confirm that our single-cell G4 profiling via scCUT&Tag can detect G4 structures in individual cells with high specificity and has sufficient resolution to distinguish cell types based on their G4 landscape.

**Figure 1:**
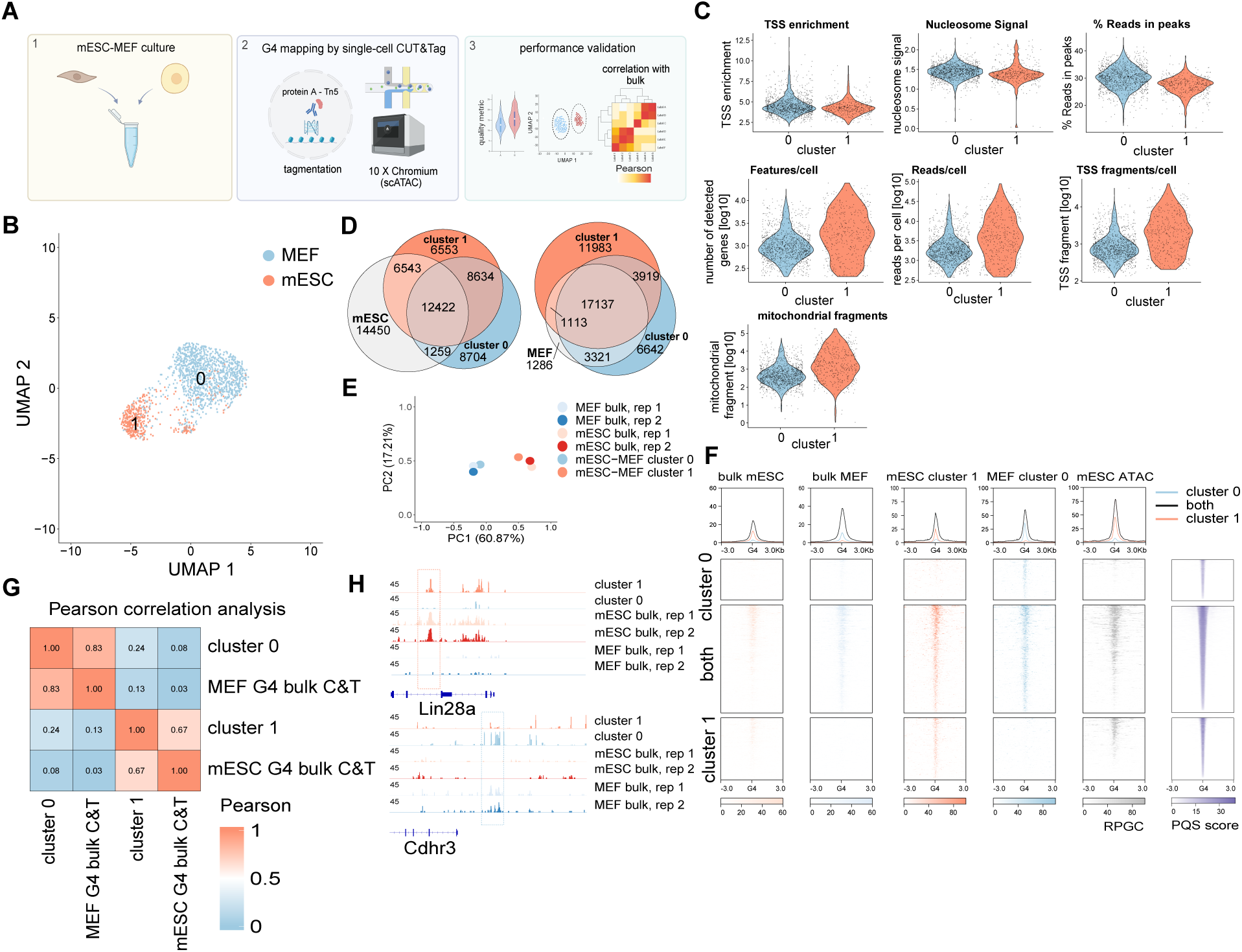
G4 profiling by scCUT&Tag is capable of separating different cell types. A.) Schematic view of the mESC-MEF G4 scCUT&Tag experimental design. A mix of two defined cell populations, mouse embryonic stem cells (mESC) and mouse embryonic fibroblasts (MEF) was used to assess the resolution of G4 scCUT&Tag. Nuclei were isolated, incubated with specific antibodies against G4s (BG4), tagmented using a protein A-Tn5 fusion, and processed following the 10x Chromium scATAC-Seq protocol. Read count matrices were processed, quality checked, normalized, and explored. B.) UMAP followed by clustering performed on the mESC-MEF G4 scCUT&Tag returns two different clusters. The resolution parameter of SNN-based clustering was set to 0.1. Cells are colored according to Seurat clusters. C.) Violin plots showing the quality measures of the different clusters. All the metrics were computed by Seurat. D.) Venn diagrams show peak overlaps among Seurat cluster 0, cluster 1, and mESC bulk G4 CUT&Tag, as well as among Seurat cluster 0, cluster 1, and MEF bulk G4 CUT&Tag. E.) PCA on the normalized signals of marker regions (output from the Seurat FindAllMarkers function) for our mESC-MEF G4 scCUT&Tag clusters shows that cluster 0 and cluster 1 group together with mESC and MEF bulk CUT&Tag replicates, respectively. F.) Heatmaps of G4 occupancy at unique and overlapping peak regions. Signals are normalized by Reads Per Genomic Content (RPGC). The PQS scores reported by the pqsfinder package indicate the G4 forming potentiality of the given region G.) Pearson correlation analysis among bulk CUT&Tag datasets and mESC-MEF Seurat clusters using the marker regions (n = 416). H.) Representative genome browser tracks of cluster 1 specific mESC G4 and cluster 0 specific MEF G4, respectively. The dashed boxes indicate the location of the unique peaks. Signals are RPGC normalized.

### Exploration of the single-cell G4 landscape in postnatal mouse brain

Having established that scCUT&Tag allowed the deconvolution of cell-type specific G4 structures in heterogeneous mixtures (**Fig 1**), we turned to the developing mouse brain, collecting P15/P25 postnatal brain cells from transgenic mice expressing Sox10:Cre/Rosa26:(CAG-LSL-EGFP(RCE)) (34). The reporter labels major brain cell types, including astrocytes (AST), mature oligodendrocytes (MOL), oligodendrocyte progenitor cells (OPC), committed oligodendrocyte precursors or newly formed oligodendrocytes (COP-NFOL), olfactory ensheathing cells (OEC), and vascular cells such as pericytes, vascular endothelial cells (VEC), and vascular and leptomeningeal cells (VLMCs). The cells without GFP were more abundant in neurons and microglia relative to the GFP+ population. We conducted G4 scCUT&Tag experiments with (sorted) or without (unsorted) fluorescent-assisted cell sorting and characterized the samples upon multi-omics integration (**Fig 2A**). First of all, we processed the GFP+ sorted population using a standard Seurat workflow (see Materials and Methods), adjusting the clustering resolution parameter to 0.8. This higher resolution allowed us to identify more clusters (**Suppl. Fig 2A**) and assess whether our G4 data - comparable in depth to other scCUT&Tag epigenetic profiles (34) - could be utilized to deconvolute the cell type composition.

**Figure 2:**
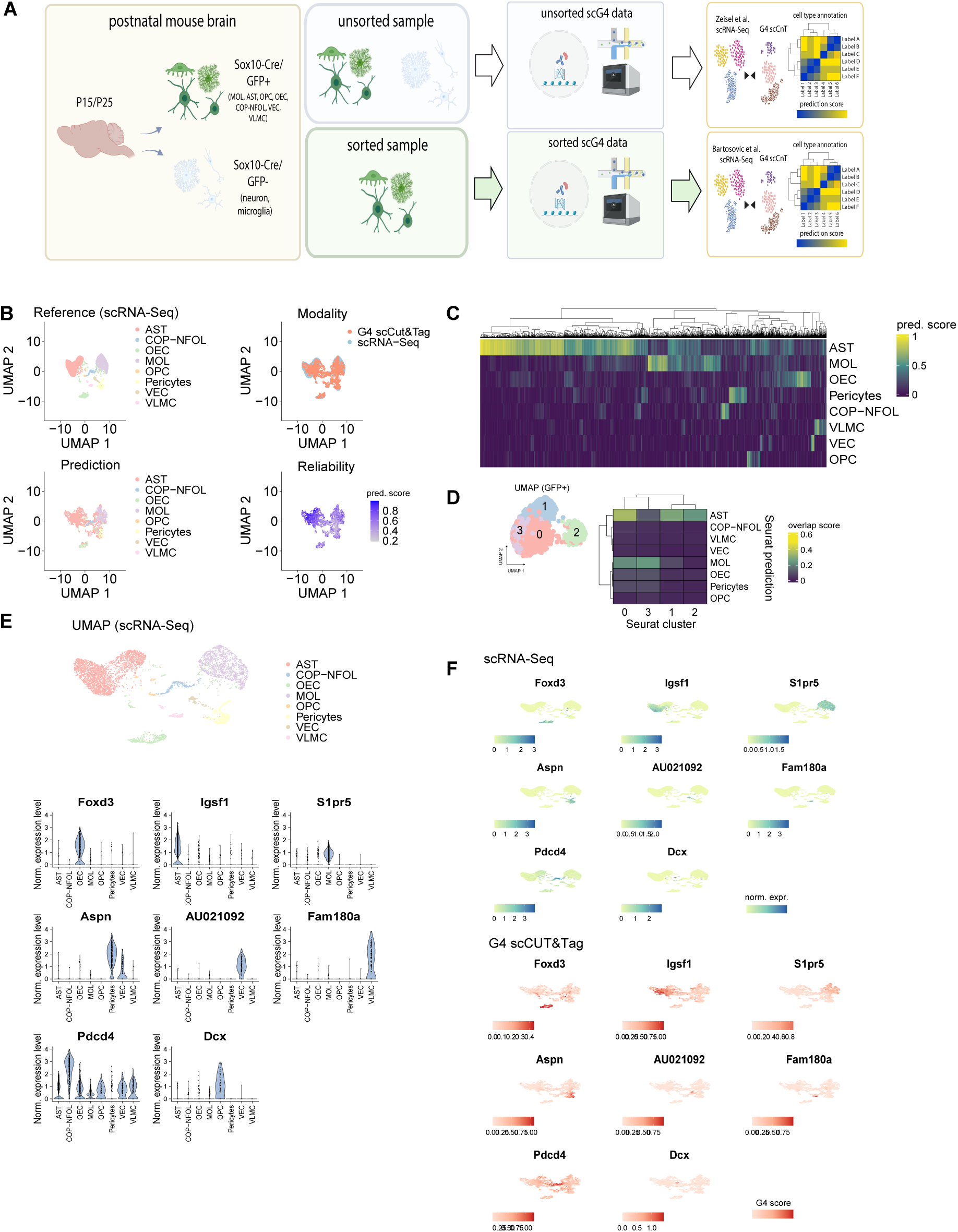
Integration of G4 scCUT&Tag data with scRNA-Seq using Canonical Correlation Analysis (CCA) A.) Schematic workflow for the study of single-cell G4 landscapes in postnatal mouse brain. B.) UMAP representations after the batch correction by CCA. Colors indicate the scRNA-Seq (34) labels upon CCA (Seurat), the data modalities, the predicted labels and the prediction scores, respectively. C.) Heatmap representing the prediction scores coming from the transferred class annotations. D.) Heatmap representing the overlap score quantifying the similarity upon label transfer between the predicted labels and GFP+ G4 scCUT&Tag clusters. E.) Output of the scRNA-Seq marker analysis. Violin plots display the distributions of marker gene expressions. F.) Co-enrichments of G4 activity scores and normalized expression signals of the defined scRNA-Seq marker genes. The corresponding scRNA-Seq clusters are highlighted in the UMAP of E.). Color scales indicate the gene activity score and normalized gene expression of G4 scCUT&Tag and scRNA-Seq, respectively. AST: astrocyte, MOL: mature oligodendrocyte, OPC: oligodendrocyte precursor, COP: committed oligodendrocyte precursor, OEC: olfactory ensheathing cells, NFOL: newly formed oligodendrocyte, VEC: vascular endothelial cells, VLMC: Vascular and leptomeningeal cell.

To annotate cell types, we used a labeled single-cell RNA-Seq (scRNA-Seq) dataset (**Suppl. Fig 2B**) from postnatal mouse brain of the same age (34) and performed horizontal data integration between the single-cell G4 and transcriptomics modalities by co-embedding the datasets using mutual nearest neighbors and canonical correlation analysis (CCA, implemented in Seurat) (**Fig 2B**). To anchor features between the datasets, we considered only the scCUT&Tag signal over the gene body and promoter region in the single-cell G4 modality. We then computed gene activity scores using the GeneActivity function (Signac package) and set these scores as the default assay for the G4 analysis. Following anchor identification across the datasets (FindTransferAnchors function, Seurat), we performed data imputation (TransferData function, Seurat) with three key settings: (1) we prioritized the top variable genes (n = 2,000) from the mean variability plot of the scRNA-Seq dataset; (2) we used the previously identified anchors between the two objects as our anchor set; and (3) we applied LSI as the dimensionality reduction technique for anchor building, given the datasets span different modalities and our goal was to capture as much biological variability as possible. This joint analysis aided the estimation of the cell type compositions of the G4 clusters. Cells with AST labels showed significantly higher prediction scores after imputation (**Fig 2B, Suppl. Fig 2C**) compared to all other labeled cells. We then assessed cluster similarity across the two modalities and calculated overlap scores (**Fig 2D**, detailed in Materials and Method and in (36)). We observed that cluster 2 of the sorted dataset consists of only ASTs, while cluster 0 and cluster 1 contain multiple cell types (mainly ASTs and MOLs). Finally, using the gene expression markers defined previously from the scRNA-Seq dataset (**Fig 2E**) and the co-embedded G4 data we asked whether transcripts and G4s show any co-enrichments at the level of cell types. The feature plots showed (**Fig 2F**) that G4 signals at the marker genes correlated with their scRNA-seq level. To sum up, we found that the GFP+ population could be deconvoluted into major cell types, demonstrating cell-type specific G4 heterogeneity, but the overall separation power of G4 signal is weaker than scRNA-seq and scATAC-seq. We also found that the expression of marker genes is accompanied by an increased G4 signal.

### Integration through reliability modeling enhances the annotation of G4 scCUT&Tag clusters

To achieve more robust cell type annotation for the GFP+ brain sample, we used scBridge (31) which is a semi-supervised method performing the integration in a heterogeneous transfer learning manner. scBridge performs annotation through a progressive learning approach, beginning with cells in the scCUT&Tag data that have the most similar profiles to scRNA-Seq cells based on higher positive correlations with gene expression. This technique allowed us to leverage not only the cell heterogeneity of the sample but also the similar enrichment patterns observed in the co-embedded data space (**Fig 2F**). To illustrate the scBridge driven integration of scRNA-Seq and G4 scCUT&Tag data, we visualized the process with UMAPs in **Fig 3A**, coloring cells by reference cell type labels from the reference scRNA-Seq, data modalities, predicted labels on the G4 scCUT&Tag layer or reliability score.

**Figure 3:**
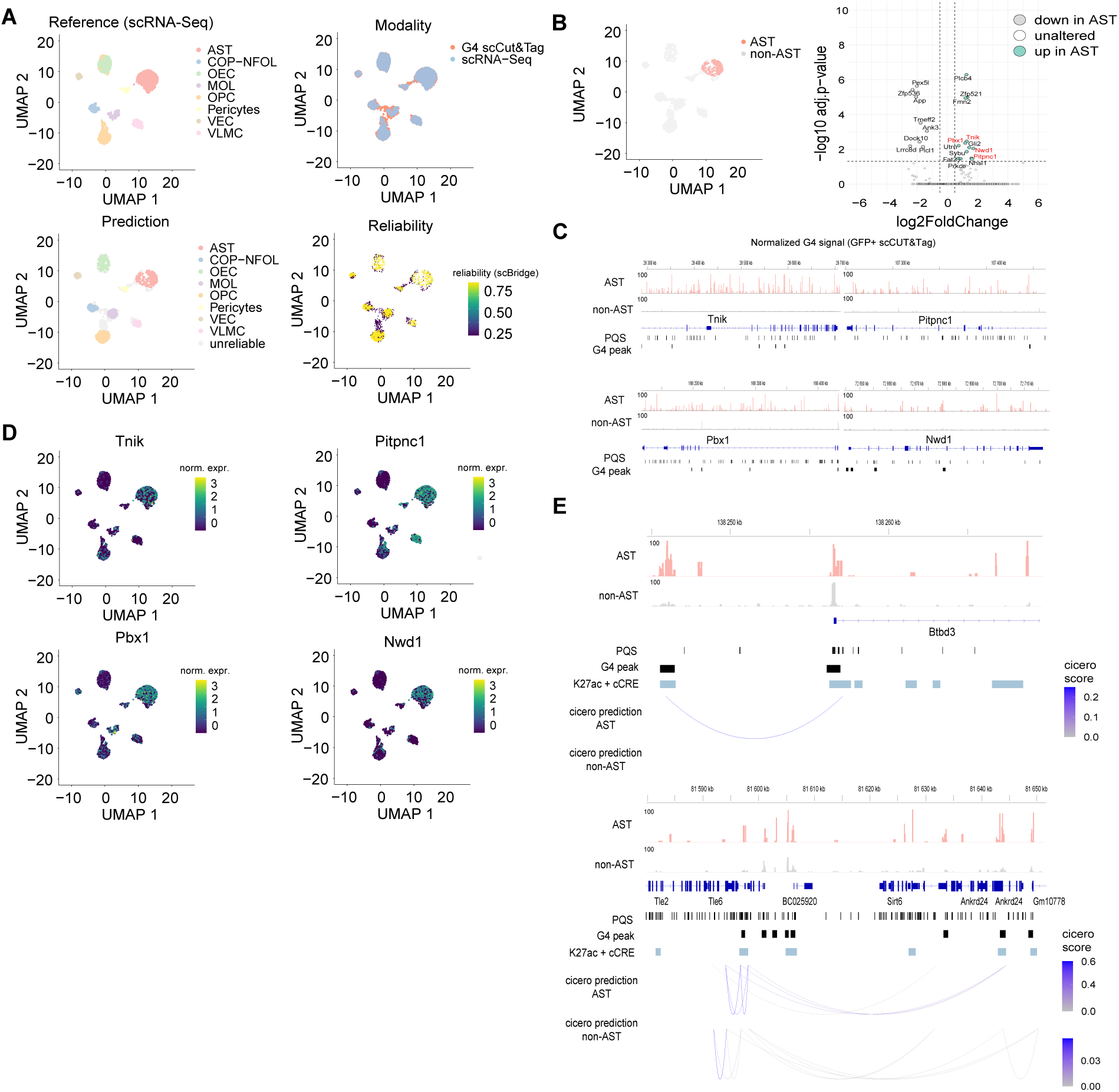
Integration through reliability modeling enhances the annotation of G4 scCUT&Tag clusters. A.) UMAP plot of the joint embedding of scRNA-seq and G4 scCUT&Tag obtained by scBridge. Cells are colored by cell types, predicted labels, integration reliability and modality, respectively. B.) Differential G-quadruplex region analysis between the predicted AST cells and cells with other, non-AST labels. Volcano plot shows the differential G4 signals in terms of activity scores. Dashed lines denote the thresholds of the differential analysis: average |log_2_ fold change| > 0.5 & adj. p-value < 0.05. For the analysis the gene activity score was considered. Red gene symbols indicate those genes that were further explored. C.) Coverage tracks show examples of genes (indicated by red color on the volcano plot) with significantly higher G4 signals in AST cells. UMAP feature plots illustrating the elevated AST specific gene expression signals (scRNA-Seq of (34)). The color from dark to bright represents the expression level from low to high. D.) Cicero predictions of AST specific G4 peaks over candidate mouse brain regulatory elements (cCREs coming from (37)). Boxes indicate the G4 peaks called by MACS2, PQS sites and regulatory elements, respectively. Only those regulatory elements were considered that showed higher H3K27ac signal than the 75th percentile of K27ac signal of all elements (H3K27ac data derives from the (34) histone modification scCut&Tag study). Coverage plots show the normalized G4 signals aggregated for the AST and non-AST clusters, respectively.

As shown, scBridge was discriminative, assigning high reliability to a substantial portion of cells (2083 cells from 2767 had reliability score > 0.9). To select a specific cell type for further characterization, we visualized Seurat prediction scores and predicted labels on the scBridge UMAPs (**Suppl. Fig 3A**) and calculated overlap scores across Seurat clusters and scBridge predictions as in **Fig 2C** (**Suppl. Fig 3B**), as well as between reliable Seurat predictions (prediction score cutoff: 0.5) and scBridge predictions (**Suppl. Fig 3C**). Because prediction scores for the AST population were most robust, we further investigated the G4 landscape of this population. First, we aimed to identify differentially enriched G4 regions associated with the predicted AST label. We aggregated the cells within the predicted AST cluster (246 cells, average scBridge reliability score: 0.98, Seurat prediction score > 0.9) and performed a differential G-quadruplex region analysis, comparing these cells to all other cells (2521 cells, hereafter non-AST) (**Fig 3B**). This analysis revealed 8 genes with significantly lower and 13 genes with significantly higher gene activity scores in AST cells (**Fig 3B**). To further explore the AST cluster, we focused on four genes (highlighted in red on the volcano plot) that showed significant G4 enrichments in AST cells and visualized their normalized G4 and gene expression signals. Coverage plots demonstrated elevated AST-specific G4 signals and numerous canonical PQS sites over the gene bodies (**Fig 3C**). Consistently, all four genes also exhibited significantly higher expression levels in the AST population (**Fig 3D**), aligning with our previous co-enrichment findings (**Fig 2C**). In summary, multi-omics integration of G4 scCUT&Tag and scRNA-Seq enabled the discovery of unique G4 patterns near astrocyte-specific genes and facilitated deconvolution of the cell-type origins within the heterogeneous mouse brain sample. Hence, cell-type specific gene expression activity appears to be a driver of cell-type specific G4s.

Lastly, we addressed the question, if G4 peaks may represent cis-regulatory regions. To explore this, we applied cicero (32) on the normalized single-cell G4 signals from the annotated GFP+ scCUT&Tag dataset. Originally developed for single-cell chromatin accessibility data, cicero is an unsupervised method that predicts cell-type-specific interactions between cis-regulatory regions and works by measuring the correlation between pairs of proximal links. Following LSI preprocessing and UMAP reduction, cicero identified 22,928 and 2,758 correlated features with co-accessibility scores above 0.2 in AST and non-AST cells, respectively. Representative examples of regions with high co-accessibility scores (>0.2) demonstrated that physically proximal G4 sites mostly overlap with H3K27-acetylated putative cis-regulatory elements (37) and that predicted loops differ between AST and non-AST cells (**Fig 3E**).

Additionally, three of our previously selected AST-specific genes also contained interacting G4 sites at H3K27-acetylated regulatory regions (**Suppl. Fig 3D**). Altogether, G4 scCUT&Tag facilitates the detection of interacting G4 sites that colocalize with cis-regulatory regions, enabling single-cell level examination of G4 interactions with putative enhancers.

### Integration of the GFP-cluster from the unsorted G4 scCUT&Tag dataset with neuronal scRNA-Seq

To further explore the G4 landscape of the developing brain, we performed an additional G4 scCUT&Tag experiment on the unsorted mouse brain tissue. Initial dimensionality reduction of the unsorted single cells (n = 3,910) and clustering with a low-resolution parameter (set to 0.1) resulted in one major cluster 0 and a smaller cluster 1. We mapped the unsorted cells (query) to the GFP+ cells (reference) and assigned labels accordingly (**Fig 4A**). Cells from cluster 0 showed significantly higher prediction scores (mean: 0.84) than those from cluster 0 (mean: 0.69) (**Fig 4B**), suggesting that cluster 0 contained most of the GFP+ population. To further characterize these clusters, we performed peak calling and identified 4,941 peaks unique to cluster 1 of the unsorted population, which were absent in both cluster 0 and the GFP+ population (**Fig 4B**). Pseudo-bulk showed a strong signal for cluster 1 specific peaks that was indeed absent in cluster 0 and the GFP+ cells. These cluster 1-specific peaks were enriched in genes associated with GO terms such as neural tube closure and primary neural tube formation, suggesting that cluster 1 consists of neuronal cell types (**Fig 4D**). For instance, genes such as *Cc2da*, *Lias*, and *Prkacb*, which are involved in primary neural tube formation, showed unique G4 signals in cluster 1 (**Fig 4E**). In order to more thoroughly dissect the cell types of cluster 1, we attempted the co-embedding of our unsorted cluster 1 cell population with scRNA-seq from an adolescent mouse brain. For the data integration, we used neuron scRNA-Seq data from the mouse brain atlas of *Zeisel et al* (38) as ground truth considering 13 distinct tissues. Indeed, the resulting cell embeddings (**Fig 4F**) displayed clear patterns corresponding to neuronal subtypes, enabling confident annotation (scBridge reliability score > 0.90) of 444 out of 963 cluster 1 cells. These annotated cells spanned various neuronal types, including those from the amygdala, thalamus, cortex, striatum, and cerebellum (**Fig 4G**), but also approximately 50% of the cells remained unannotated. This might be due to the limitations of the label transfer or the presence of cell types not represented in the scRNA-seq reference, such as non-neuronal populations.

**Figure 4:**
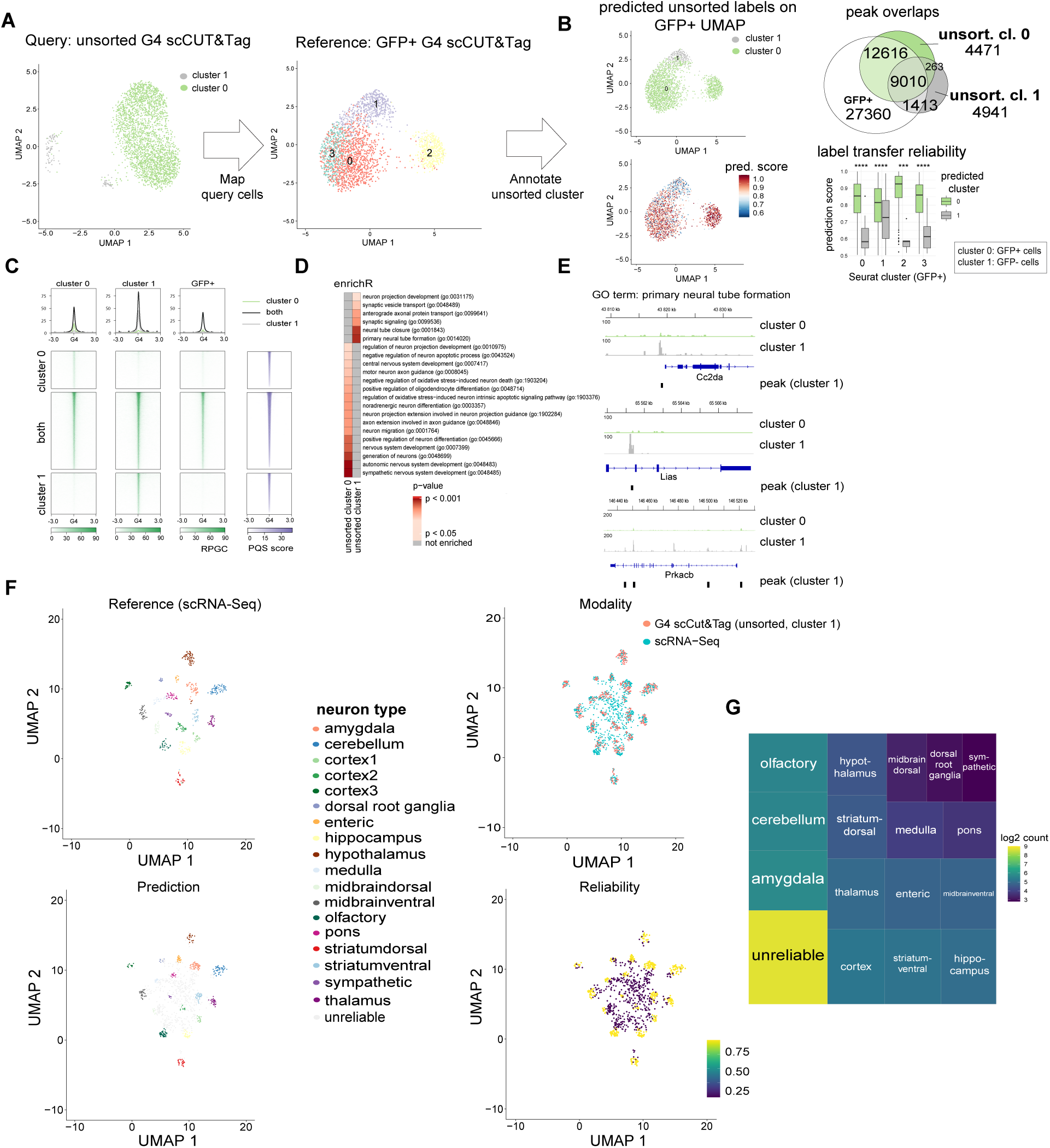
Integration of the GFP-cluster from the unsorted G4 scCUT&Tag dataset with neuronal scRNA-seq aids the identification of neuronal cell types. A.) Projection of the unsorted data as a query (containing GFP- and GFP+ fractions) onto the GFP+ dataset as a reference. B.) UMAPs show the mapped query labels and the prediction scores upon the label transfer. Venn diagram indicates the peak overlaps between the GFP+ dataset and unsorted clusters. Boxplot shows the distribution of the prediction scores between the unsorted clusters at the level of GFP+ Seurat clusters. C.) Heatmaps of G4 occupancy at unique and overlapping peak regions. Signals are normalized by RPGC. The PQS scores reported by the pqsfinder package indicate the G4 forming potentiality of the given region D.) enrichR gene ontology analysis on the cluster-specific peak sets. Heatmap indicates the CNS-related GO terms and the corresponding p-values. E.) Genome viewer tracks display representative cluster 1 specific G4 peaks. The genes are related to the GO term primary neural tube formation (GO:0014020). F.) UMAP visualizations of the joint embedding of *Zeisel et al.* scRNA-seq data and the cluster 1 of the unsorted dataset. Cells are colored according to neuron types, data modality, predicted labels, and integration reliability, respectively. G.) Treemap quantifying the predicted neuron types. The color scale represents the log_2_-transformed count of each neuron type inferred from label imputation.

## Discussion

Emerging single-cell technologies offer unprecedented opportunities to explore cellular heterogeneity and to uncover the underlying regulatory epigenetic processes (38) (34) (35) (39) (40). G4 scCUT&Tag is a previously deployed technique (28) that enables the profiling of G4s at the single-cell level and allows us to investigate cell-to-cell variation of this DNA secondary structure. To assess the performance of our experimental setup, we first compared the processed single-cell G4 data with bulk G4 CUT&Tag datasets obtained from a centrifugation-based protocol, which outperformed our earlier bead-based CUT&Tag workflow. (**Suppl. Fig 1A-D**). The normalized read coverage tracks from aggregated single cells showed strong correlation with bulk G4 CUT&Tag data for both mESC (Pearson > 0.6) and MEF (Pearson > 0.8), confirming that our single-cell data accurately recapitulates the bulk G4 CUT&Tag observations. The library complexity of our mixed samples, characterized by median high-quality fragment counts of 3,434, 1,983, and 1,511 per cell, and FRiP scores of 28.3%, 13.5%, and 16.3% for the mESC-MEF, sorted, and unsorted mouse datasets, respectively, was comparable to a previous G4 scCUT&Tag study on a mixture of two human cell lines, which reported 1,226 fragments with a FRiP score of 26% for U2OS and 819 fragments with a FRiP score of 10.5% for MCF7 (28). We also demonstrated through experiments on the sorted and unsorted postnatal mouse brain samples that G4 scCUT&Tag enables the study of G4 signatures in the developing mouse brain. Using multi-omics integration, we successfully deconvoluted the cell type compositions of the samples through two distinct approaches: (i) a linear combination of features via CCA and (ii) heterogeneous transfer learning with scBridge, a deep learning-based tool. In addition to identifying distinct cell populations de novo, we observed unique G4 enrichments across different clusters. For instance, we detected significant G4 enrichment at the promoter region of *Klk6* in the cluster 0 of the unsorted sample, a gene highly expressed in oligodendrocytes in the normal CNS (41) (42), and in reactive astrocytes and microglia under inflammation or trauma (43) (44) (**Suppl. Fig 3E**). Although, upon annotation of the sorted brain sample, only the predicted astrocyte (AST) cluster exhibited G4 enrichment at the *Klk6* promoter (AST vs. non-AST average log2 fold change: 0.85, not statistically significant). Additional examples of cell type-specific G4s include the genes *Gfap* (**Suppl. Fig 3F**) and *Idh1* (**Suppl. Fig 3G**). *Gfap*, an intermediate filament protein, is characteristic of both mature and developing astrocytes in the CNS (45) (46), while *Idh1* is predominantly expressed in oligodendrocyte-lineage cells (47) and is frequently mutated in adult gliomas (48) (49). Furthermore, we observed cluster-specific G4 peaks over genes responsible for primary neural tube formation (**Fig 4E**) and predicted unique G4 formations at potential cis-regulatory elements specifically in the AST population (**Fig 3E**). Overall, identifying unique G4 peaks at genomic regions such as promoters, gene bodies or potential enhancers enables the exploration of G4 dynamics in relation to cell type-specific epigenetic and transcriptional changes in various biological processes (50) (51) (19). We sought to demonstrate the resolution and versatility of G4 scCUT&Tag data through various downstream analyses, including multi-omics data integration and regulatory region linkage. To annotate the exact cellular identities of the GFP+ population and assess the integrability of our single-cell data, we aligned our sorted G4 scCUT&Tag assay with a scRNA-Seq dataset from postnatal mouse brain tissue of the same age (34). Using canonical correlation analysis with correlation alignment, we anchored these two independent datasets, projecting them into a shared feature space. The resulting co-embedded representation revealed co-enrichment between aggregated G4 signals and transcriptional levels of key marker genes, suggesting an interplay between G4 structures and active transcription. This supports recent findings that G4s are predominantly located at gene promoters of transcriptionally active genes, where they act predominantly as transcriptional enhancers (52) (53) (50) (54), rather than as roadblocks or repressors (55) (20). However, determining the transcriptional causality of endogenous G4s remains challenging and would benefit from paired rather than unpaired multi-omics integration. The implementation of multi-omics approaches, such as simultaneous measurement of G4 signal and gene expression within the same cell, would provide deeper insights into the potential causal relationships between G4 structures and gene expression.

Another goal of our integration was to show that, similar to histone modification scCUT&Tag of *Bartosovic et al.*, G4 scCUT&Tag enables cell type annotation upon the scRNA-Seq integration. Notably, deconvolution using CCA was less straightforward here than in previous studies involving scATAC-Seq (37) (56) or histone modification scCUT&Tag (34), as reflected by the prediction scores from the anchoring process (**Suppl. Fig 3E**). These observations suggested that G4 signal did not merely follow cell-type specific DNA accessibility or histone PTM patterns, but were concentrated on a relatively small number of promoters and cis-regulatory elements while the majority of G4 resided in house-keeping and broadly expressed genes. To mitigate potential misclassifications, we conservatively selected cells with relatively high prediction scores for downstream analyses, retaining, for example, only cells with prediction scores above 0.9 for AST characterization. However, CCA does not account for sample heterogeneity or the non-linear nature of high-dimensional single-cell multi-omics data. To address this, we tested a recent deep learning tool, scBridge, and re-integrated the datasets with a strict reliability threshold. By combining reliable predictions from both CCA and scBridge, we selected the predicted AST cluster to characterize its G4 landscape, focusing on differential G-quadruplex sites (based on gene activity scores) and transcriptional levels of the genes with differential G4 coverage. As we expected, many genes with elevated G4 signals in the AST cluster showed AST-specific transcriptional patterns. Exceptions were *Gli2* (AST vs. non-AST average log_2_ fold change: 1.5, adj. p-value ***) and *Zfp521* (AST vs. non-AST average log_2_ fold change: 1.1, adj. p-value ***), that showed significantly higher G4 fold changes in the AST population but lacked expression in any cluster. We focused on the AST cluster, as it returned the highest annotation accuracy based on the overlap score matrix (**Suppl. Fig 3C**). Similar to the sorted data, we annotated the unsorted sample to estimate the neuronal composition of the cluster enriched in neuronal cell types (**Fig 4E**). Taken together, integrating G4 scCUT&Tag with scRNA-Seq revealed AST- and neuron-specific G4 profiles in heterogeneous brain samples, suggesting that CNS cell type composition is reflected at the G4 level.

Finally, we studied the G4s beyond their local context and addressed the question, whether we are able to capture linearly distant but spatially interacting G4 sites (57) (58) (59). Using cicero, we successfully identified cluster-specific G4 co-occurrences, some of which were located within potentially active regulatory elements marked by high H3K27ac levels (37) (**Suppl. Fig 3D**) (**Fig 3C**). Consequently, G4 scCUT&Tag allows to examine the dynamics and cell type specificity of G4s beyond its local context and could consider cell type specific roles in long-range enhancer-promoter interactions. Studying single-cell G4 dynamics at both chromosome conformation and cis-regulatory levels may be particularly relevant to the G4 and 3D chromatin fields, given the possible roles of G4s in forming enhancer-promoter loops (60) (61) (62) (63) (64). While G4 ChIP-Seq (19) and CUT&Tag (65) (33) (66) have generated maps of folded G4 sites across various cell types, states, and diseases, these techniques have either lacked access to sub-population level insights or have not examined them in depth (28). We anticipate that G4 scCUT&Tag, combined with computational tools commonly used in single-cell omics workflows, such as scRNA-Seq and scATAC-Seq, will provide a versatile framework for analyzing the dynamics and cell type specificity of these secondary DNA structures. However, in its current form, this technique does not reveal sub-populations in an unsupervised manner (**Suppl. Fig 2A**). In our study, differentiating cell types in the heterogeneous sample required integration with scRNA-Seq data. Extending G4 scCUT&Tag to a paired multi-omics approach - enabling simultaneous measurement of scCUT&Tag and RNA-Seq signals within the same cell - could shed light on causal relationships between G4 structures and gene expression. Additionally, we anticipate that technological advancements, particularly in increasing the number of unique fragments per cell, will further enhance our ability to resolve G4s in their endogenous cellular contexts.

## Materials and methods

### Bulk G-quadruplex CUT&Tag with centrifugation

Bulk G-quadruplex CUT&Tag experiments were performed as in Lyu et al. (33), but without Concanavaline A beads to better maintain the integrity of the nuclei. All the centrifugation steps in the protocol were performed at 600g or 1700rpm at 4℃ with swing bucket centrifuge. 200,000 cells were harvested by centrifugation, resuspended in wash buffer (20 mM HEPES pH 7.5, 150 mM NaCl, 0.5 mM spermidine) and pelleted again, incubated in 200ul of antibody binding buffer (wash buffer with 2 mM EDTA, 0.01% NP-40, 0.05% digitonin and 1% BSA) containing 2ug of BG4 antibody (Addgene #55756) with rotation overnight at 4℃. The next day, cells were washed with 200ul of dig-wash buffer (wash buffer with 0.01% NP-40, 0.05% digitonin and 1% BSA) three times by centrifugation at 600g or 1700rpm for 4min with swing bucket centrifuge. After washing, cells were incubated with 1:100 dilution of anti-FLAG antibody (Sigma, F1804) for one hour at room temperature with rotation. Cells were washed with 200ul of dig-wash buffer to remove unbound anti-FLAG antibodies. Anti-FLAG treated cells were incubated with 1:100 dilution of anti-mouse antibody (Sigma, M7023) in a dig-wash buffer for one hour at room temperature, followed by washing with 200ul of dig-wash buffer three times. After antibody incubation, cells were incubated with 1:200 dilution of assembled pA-Tn5 in 200ul of dig-300 buffer (20 mM HEPES pH 7.5, 300 mM NaCl, 0.5 mM spermidine, 0.01% NP-40, 0.05% digitonin and 1% BSA) for 1h at room temperature. After pA-Tn5 binding, cells were washed with 200ul of dig-300 buffer three times. Afterwards, pA-Tn5 bound cells were incubated in 200ul of tagmentation buffer (dig-300 buffer with 10mM MgCl2) at 37℃ for 1h without shaking. After tagmentation, genomic DNA was extracted and purified with DNA Clean & Concentrator-5 (Zymo research, D4013). To generate the G4 library, purified genomic DNA was amplified with the universal i5 primer and barcoded i7 primer using NEBNext Ultra II Q5 Master Mix (NEB, M0544). The library PCR products were cleaned up with Agencourt AMPure XP beads (Beckman Coulter, A63881) and sequenced on an Illumina Nextseq 2000 instrument.

### pA-Tn5 loading

Mosaic-end adapter A (Tn5ME-A, TCGTCGGCAGCGTCAGATGTGTATAAGAGACAG) and Mosaic-end adapter B (Tn5ME-B, GTCTCGTGGGCTCGGAGATGTGTATAAGAGACAG) oligonucleotides were each annealed with Mosaic-end reverse oligonucleotides (Tn5MErev, 5′- (phos)CTGTCTCTTATACACATCT-3′). To anneal, the oligonucleotides were diluted to 200 µM. Each pair of oligos, Tn5MErev/Tn5ME-A and Tn5MErev/Tn5ME-B, was mixed and annealed separately. The loading of pA–Tn5 with the oligonucleotides was performed by incubating 20ul of pA-Tn5 (5.5uM) with 2.2ul of annealed Tn5MErev/Tn5ME-A (100uM) and 2.2ul of annealed Tn5MErev/Tn5ME-B (100uM) at room temperature for 1h.

### Cell line culture

Mouse embryonic stem cells were cultured feeder-free in 0.1% gelatin-coated flasks (Sigma, G1890) under standard conditions (5% CO2, 90% humidity, 37°C) in KnockOut DMEM (LifeTechnologies, 10829018), 2 mM alanyl-glutamine (Sigma, G8541), 0.1 mM non-essential amino acids (Sigma, M7145), 15% fetal bovine serum (FBS) (Sigma, F7524), 0.1 mM β-mercaptoethanol (Sigma, M3148), ESGRO Leukemia Inhibitory Factor (LIF) (Millipore, ESG1107), 1 μM PD0325901 (PZ0162-25MG) and 3 μM CHIR99021 (SML1046-25MG). NIH-3T3 cells were cultured in DMEM (Gibco, 31966047) supplemented with 10% fetal bovine serum (FBS; Gibco, 26140079) and 1× penicillin/streptomycin (Sigma, P4333).

### Animals

The Sox10:Cre/Rosa26:(CAG-LSL-EGFP) mice (RCE) mouse line used in this study was previously described by Bartosovic et al. (34). All animals were free from mouse viral pathogens, ectoparasites and endoparasites, and mouse bacteria pathogens. Mice were kept with the following light/dark cycle: dawn 6:00–7:00, daylight 7:00–18:00, dusk 18:00–19:00, night 19:00– 6:00 and housed to a maximum number of five per cage in individually ventilated cages (IVC sealsafe GM500, tecniplast). General housing parameters such as relative humidity, temperature, and ventilation follow the European convention for the protection of vertebrate animals used for experimental and other scientific purposes treaty ETS 123, Strasbourg 18.03.1996/01.01.1991. In brief, consistent relative air humidity of 55 ± 10%, 22 °C and the air quality is controlled with the use of standalone air-handling units supplemented with HEPA-filtrated air. Monitoring of husbandry parameters is done using ScanClime (Scanbur) units. Cages contained hardwood bedding (TAPVEI, Estonia), nesting material, shredded paper, gnawing sticks, and card box shelter (Scanbur). The mice received regular chow diet (either R70 diet or R34, Lantmännen Lantbruk). Water was provided by using a water bottle, which was changed weekly. Cages were changed every other week. All cage changes were done in a laminar air-flow cabinet. Facility personnel wore dedicated scrubs, socks, and shoes. Respiratory masks were used when working outside of the laminar air-flow cabinet. Animals were sacrificed at juvenile stages (P19) and both sexes were included in the study. All experimental procedures on animals were performed following the European directive 2010/63/EU, local Swedish directive L150/SJVFS/2019:9, Saknr L150 and Karolinska Institutet complementary guidelines for procurement and use of laboratory animals, Dnr. 1937/03–640. The procedures described were approved by the local committee for ethical experiments on laboratory animals in Sweden (Stockholms Norra Djurförsöksetiska nämnd), license number 144/16, 1995_2019 and 7029/2020.

### Tissue dissociation

Mice were sacrificed, perfused with 1× PBS, pH 7.4 and the brain was removed. The brain was dissociated into single-cell suspension using Neural Tissue Dissociation Kit P (Miltenyi Biotec, 130-092-628) according to the manufacturer’s protocol. For mice at P25 stage, myelin was removed using debris removal solution (Miltenyi Biotec, 130-109-398) according to the manufacturer’s instructions. Single-cell suspension was filtered through a 50 µm cell strainer and briefly stained with 1:5,000 diluted DAPI (1 mg ml−1) to assess cell viability. For FACS, cells were resuspended in 1× PBS supplemented with 1% BSA and 2 mM EDTA and GFP+ cells were sorted using fluorescent-assisted cell sorting (FACS Aria III or FACS Aria Fusion) directly into 500 μl of antibody buffer.

### Single cell G-quadruplex CUT&Tag

Single cell workflow with 10x platform was performed as in Bartosovic et al. (34). For mESC-MEF, 100,000 mESC and 100,000 NIH-3T3 cells were mixed before G4 antibody incubation. All incubation and washes are the same as in the case of bulk G4 CUT&Tag. After tagmentation, the sample was centrifuged for 3 min at 300g and resuspended in 15–25 μl of 1× PBS + 1% BSA and processed using 10x Chromium Single Cell ATAC-Seq kit. Final library amplification was performed according to the Chromium Single Cell ATAC Library kit manual sequenced on an Illumina Nextseq 2000 instrument.

### Bulk CUT&Tag analysis

Bulk CUT&Tag fastq files were processed through the nf-core/atacseq pipeline (v1.2.1). Essentially, reads were aligned to the mm10 reference genome with BWA (v0.7.17-r1188), deduplicated with Picard (v2.23.1), peaks were called with MACS2 (v2.2.7.1). deepTools (v3.5.2) (72) bamCoverage was used to produce bigwig files using RPGC (1x Genome coverage) normalization.

### scCUT&Tag data processing and explorative analysis

*Fastq alignment.* One library of the mESC-MEF sample and one-one library for the sorted (GFP+ cells) or unsorted (GFP- and GFP+ cells) postnatal mouse brain cells of Sox10:Cre/Rosa26:(CAG-LSL-EGFP) animals on P21 were prepared and sequenced on a Nextseq 2000 instrument. Fastq files were initially processed using the count function of 10X Genomics Cell Ranger ATAC software package (cellranger-atac-2.0.0, https://github.com/10XGenomics/cellranger-atac) with standard parameters. Reads were aligned to the mm10 genome reference. Cell Ranger ATAC identified 3,434, 1,983 and 1,511 median high-quality fragments per cell in the mESC-MEF, sorted and unsorted mouse brain samples, respectively, and 28.3%, 13.5% and 16.3% of these fragments, respectively, were overlapped with peaks computed by Cell Ranger ATAC.

*Quality control.* Processing of Cell Ranger ATAC H5 files and the subsequent single-cell analysis workflow were done in R using packages Seurat (v5.0.3) and Signac (v1.13) (30). Chromatin assay object was created from features detected at least in 10 cells and from cells including at least 200 features. Doublets (**Suppl. Fig 1E**) were eliminated by the R package called DoubletFinder (67) (https://github.com/chris-mcginnis-ucsf/DoubletFinder) with parameters pN = 0.25, pK = 0.09, nExp = 0.04, principal components 1:10. Quality measure plots, such as the one in Fig 1C, were visually inspected at the level of Seurat clusters and used for quality assurance. FRiP of the bulk CUT&Tag experiments was calculated with Subread (v2.0.3) featureCounts (68). *Dimensionality reduction and clustering.* After quality control and filtering, the read count matrix was transformed by term frequency inverse document frequency (TF-IDF) transformation followed by partial singular value decomposition. UMAP (69) preceded by latent semantic indexing (LSI) was run on the 2:30 dimensions, omitting the first component that highly correlates with sequencing depth. For the optimization of UMAP parameters we tested the number of neighbors (step between 5 and 50 by 2) and minimum distance (tweaked 25 values between 0.001 and 0.5). All cluster definition was performed by FindClusters of Seurat, so that the algorithm was set to smart local moving (SLM) (70) and the optimal resolution and granularity of clustering was tuned by the resolution parameter (step between 0.1 and 1 by 0.1). Eventually, for identification of cell origin in the mESC-MEF and in the unsorted mouse samples, our final parameter set was next: resolution 0.1, minimum distance 0.4, number of neighbors 30, SLM algorithm. For the sorted mouse sample, the parameter set remained unchanged, except for the resolution, which was adjusted to 0.8.

*Peak calling and bam computation.* Cluster-specific peaks were retrieved by Signac’s CallPeaks function using a simple MACS2 (71) peak calling (with the default parameters of CallPeaks) and with group.by = “seurat_clusters” to call peaks on each cluster separately. Generating bam files from each Seurat cluster was performed by the filterbarcodes function of the package called sinto (https://timoast.github.io/sinto/). The retrieved bam files were converted to RPGC normalized bigwig files by bamCoverage of deepTools.

*Finding marker regions.* The term marker is used to refer to a significant differentially G-quadruplex region between any cluster to all other clusters. Marker regions for each cluster were calculated using the FindAllMarkers function from the Seurat package. Marker detection was performed using the logistic regression model of the function, as suggested by (73), and added the total number of fragments as a latent variable to mitigate the effect of differential sequencing depth on the result. In the comparison of AST and non-AST cells (**Fig 3**) the gene activity matrix (generated by the Signac’s GeneActivity function that aggregates the counts over the gene body and promoter region) was set as input, so that the gene activity scores were generated by default base extension arguments (extend.upstream = 2000, extend.downstream = 0).

*Cicero*. Interplays across G4 sites were analyzed by the R package called cicero (32). G4 peaks were set as input with UMAP reduction and LSI preprocessing step. For finding cis-G4 networks, the “cutoff override” parameter was set to 0.2. Predictions were quantified by the co-accessibility score showing how correlated two G4 sites are in the cells. Visualization of the sites was made by the plot_connections function of the package and IGV.

*Putative G4 sequence score (PQS score).* PQS scores, used to quantify the G-quadruplex-forming potential of the peak regions, were calculated using the pqsfinder R package (74).

*Enrichment analysis.* Enrichment analysis for gene ontology was done by the R package enrichR (v3.2) (75) using the “GO_Biological_Process_2023” database. For visualization, we retained only the GO terms connected to the CNS.

*Visualizations.* IGV (Integrative Genomics Viewer) (76) tool was used for the visualization of bigwig tracks and bed files. Profile plots, heatmaps and fingerprint plot were generated by deepTools. All further R visualization was done by ggplot2. (77)

### scRNA-Seq data processing

*Bartosovic et al. scRNA-Seq.* scRNA-Seq reads derived from GFP+ cells were aligned to the mm10 reference genome using Cell Ranger. The resulting read count matrix was processed with Seurat, including log normalization to 10,000 counts (using the NormalizeData function), identification of variable features (FindVariableFeatures function with selection.method = “vst”), scaling (ScaleData function), and dimensionality reduction via PCA (RunPCA function) and UMAP (RunUMAP function). Gene expression markers were calculated by the FindAllMarkers function with min.pct = 0.5.

*Zeisel et al. scRNA-Seq.* Loom files corresponding to neurons from adolescent mouse brains were downloaded in Level 2 format from the Sten Linnarsson Mouse Brain Atlas (38) (see Data Availability). These files were further handled by the loomR package (https://github.com/mojaveazure/loomR). Afterwards, the neuronal data were processed using the same methods applied to the *Bartosovic et al.* scRNA-Seq dataset.

### scCUT&Tag integration with scRNA-Seq data

*Integration using CCA of Seurat*. Identification of anchor correspondences between the G4 scCUT&Tag of sorted mouse brain sample and *Bartosovic et al.* scRNA-Seq dataset (both derived from Sox10:Cre/Rosa26:(CAG-LSL-EGFP) mice on P21) was identified using the FindTransferAnchors function of Seurat using our scCUT&Tag data as query and the labeled scRNA-Seq data as reference. In the case of scCUT&Tag query the gene activity score assay was used for the integration according to the G4 enrichment of gene promoter and gene body as previously described (30). The computed anchor points represent two cells (with one cell from each dataset) that originate from a common biological state. Prior to anchor detection the datasets were placed in a shared low-dimensional space by CCA (29). Following the dimensional reduction nearest neighbors were determined by K-nearest neighbors and mutual nearest neighbors algorithms. Imputation was conducted by the TransferData function of Seurat using 50 dimensions and was applied only on the highly variable features of the scRNA-Seq dataset retrieved by VariableFeatures function (number of features = 2000). Weight reduction was set to the previously computed LSI representation of the G4 scCUT&Tag module.

*Integration using scBridge.* For scBridge, AnnData objects were created using the Scanpy Python package (v1.9.8) (78), derived from the gene activity matrix of scCUT&Tag and the normalized count matrix of the corresponding scRNA-Seq dataset (*Bartosovic et al.* for sorted data and *Zeisel et al.* scRNA-Seq for unsorted data). Then, the objects were converted to .h5ad files. Only those genes were kept that can be found in both assays. The .h5ad files were fed into the scBridge pipeline, so that the reliability threshold was set to 0.8. We visualized the joint embedding by UMAP. The scBridge experiment was conducted on Rocky Linux 8 OS using 4x NVIDIA Tesla T4 GPU with CUDA 12.4.

*Mapping query cells to reference.* The unsorted G4 scCUT&Tag data were projected onto the sorted UMAP using the MapQuery wrapper function in Seurat. Anchor sets were determined with the FindTransferAnchors function, using CCA reduction with prioritizing gene activity assays. For the visualizations both the reference.reduction and reduction.model parameters were set to ‘umap’. The mapping efficiency was quantified by the corresponding prediction scores of the TransferData function.

### Visualization and quantification of integration efficiency

*Prediction score of Seurat.* Following classification of cells between modalities based on the scRNA-Seq annotations, TransferData returns a prediction score assay. The calculation of prediction scores is detailed in *Stuart et al*. (30). This metric was used to quantify the label transferring efficiency across the different clusters.

*Overlap score.* To quantify the similarity between cell clusters from the two modalities, we calculated an overlapping score as the sum of the minimum proportion of cells in each cluster that overlapped within each co-embedding cluster (36) (37). Cluster overlaps varied from 0 to 1 and were visualized as a heatmap.

*Reliability score of scBridge.* The reliability score calculation and the reliability modeling of the cells are detailed in the original scBridge paper (31).

### Statistics

Plot statistics were calculated using the R package ggpubr (v0.6.0). The exact statistical tests are indicated in the figure legends. Significance of marker regions reported was tested using logistic regression with likelihood ratio test implemented in package Seurat with function FindAllMarkers. Cicero predictions with an override cutoff > 0.2 were selected for downstream analysis. For the quantification of overlaps Jaccard index was used. Venn diagrams were calculated and visualized by the eulerr package (https://CRAN.R-project.org/package=eulerr).

## Data availability

The G4 scCUT&Tag data generated for this study have been deposited at the Gene Expression Omnibus under accession code: GSE291468. Datasets from prior studies that were analyzed here: GSE163532 (GFP+ postnatal mouse brain P21 scRNA-Seq), GSE151058 (mESC ATAC-Seq). *Li et al.* adult mouse brain cCREs are available here: http://catlas.org/mousebrain. Neuronal scRNA-Seq data (level 2, loom files) were derived from the Mouse Brain Atlas of the Sten Linnarsson lab (http://mousebrain.org/).

## Code availability

https://github.com/heteyszabolcs/G4_scCUTnTag

## Acknowledgements

Fig 1A, Fig 2A were generated with a paid subscription of BioRender (www.biorender.com). We thank the National Academic Infrastructure for Supercomputing in Sweden (NAISS) funded by the Swedish Research Council through grant agreement no. 2022-06725 and their support teams. The supercomputing cluster UPPMAX was used under projects NAISS 2023/6-19, NAISS 2023/22-84, SNIC 2022/6-14, NAISS 2024/22-108. The PDC Center for High Performance Computing, KTH Royal Institute of Technology, Sweden, was used under projects NAISS 2025-22-67 and NAISS 2025-6-32. We acknowledge funding from Vetenskapsrådet, Sweden (2020–04313) (to S.J.E.); KI StratRegen Junior Project Grant (to S.J.E.); The Ming Wai Lau Center for Reparative Medicine, Sweden (to S.J.E.); Stiftelsen för Strategisk Forskning (FFL7) (to S.J.E.); The Knut och Alice Wallenbergs Stiftelse, Sweden (to S.J.E.).

## SUPPLEMENTARY FIGURES

**Supplementary Fig 1.**
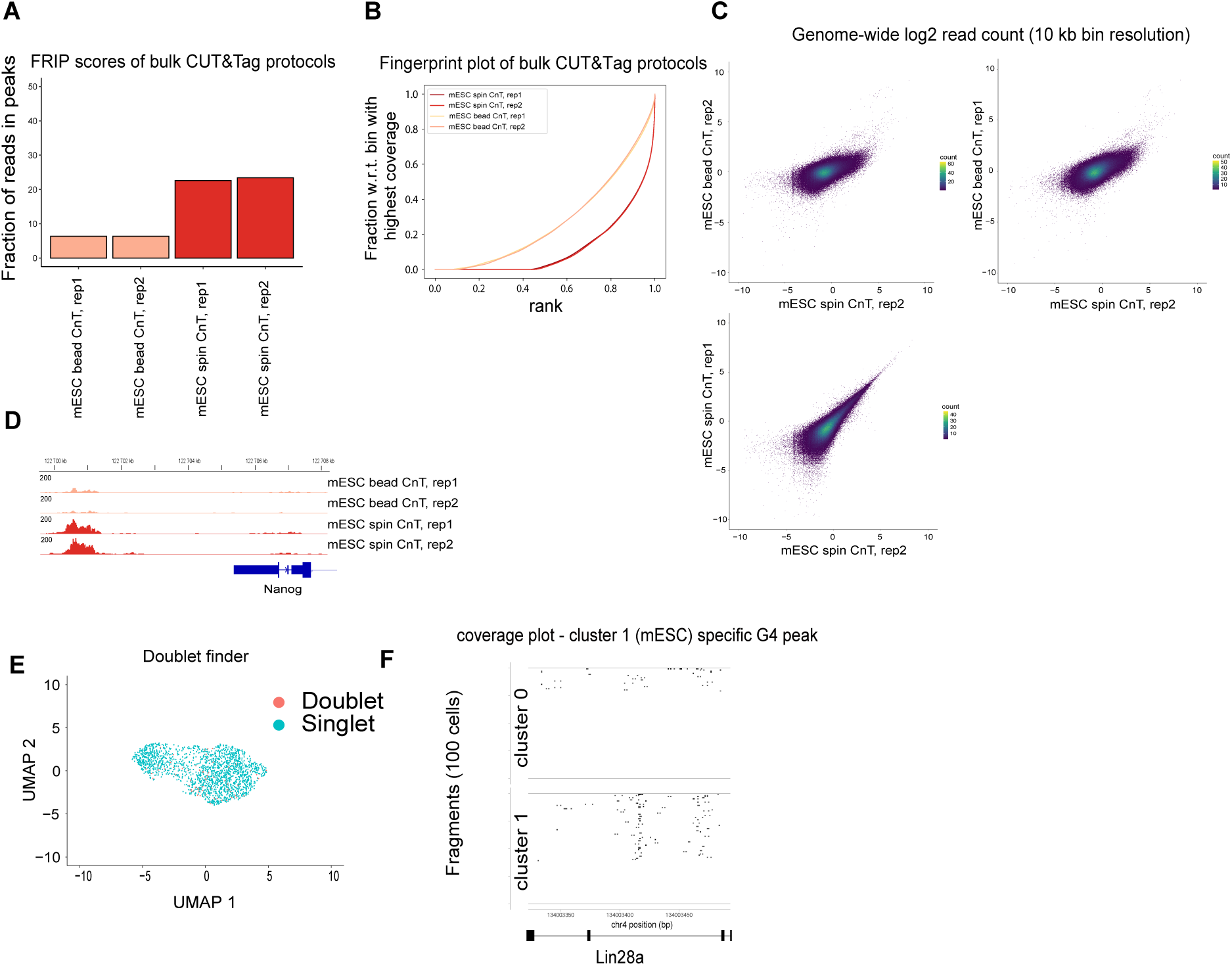
Spin-based G4 CUT&Tag outperforms bead-based protocol. A) Differences in FRiP scores between bead- and spin-based bulk CUT&Tag protocols. B) Fingerprint plot (computed using deepTools) showing cumulative read coverage differences between the two bulk CUT&Tag protocols. C) Distribution of log_2_ read coverage across 10 kb genomic windows. D) Representative G4 landscape near the *Nanog* gene, illustrating signal-to-noise differences between bead-and spin-based protocols. E.) UMAP of mESC-MEF G4 scCUT&Tag with the indication of identified doublets (by package DoubletFinder). F.) Coverage plot of representative mESC cluster-specific G4 peak over the gene *Lin28a*.

**Supplementary Fig 2.**
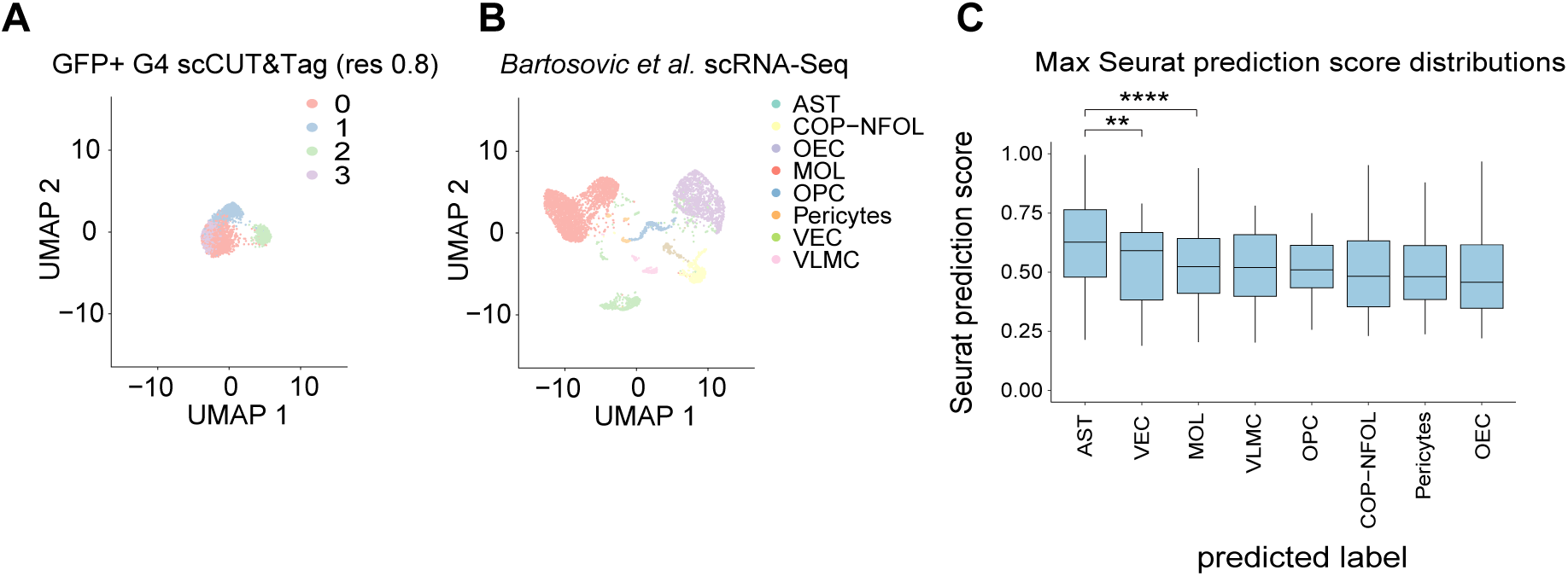
Seurat clustering of G4 CUT&Tag and scRNA-seq data. A.) UMAP representation of the GFP+ sorted G4 scCUT&Tag before the integration made by CCA (Seurat). For clustering resolution parameter was set to 0.8 with the SLM algorithm. B.) UMAP of the labeled, postnatal mouse brain scRNA-Seq dataset before the CCA. C.) Distributions of the maximum Seurat prediction scores per cell across the different cell types. P-values are calculated from two-tailed, Wilcoxon rank-sum tests.

**Supplementary Fig 3.**
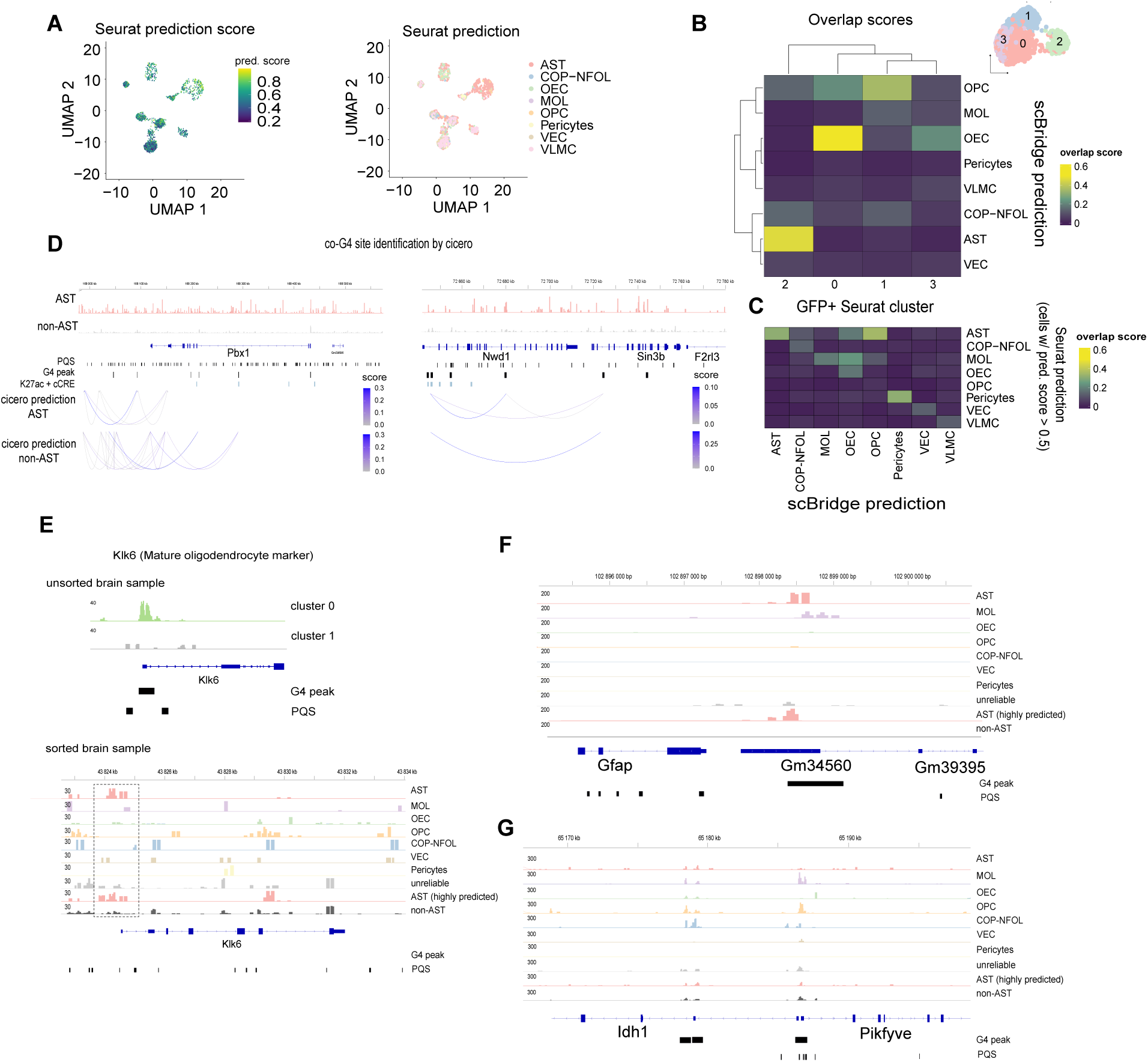
Seurat clustering of G4 CUT&Tag and scRNA-seq data. A.) UMAP plot of the joint embedding of scRNA-seq and G4 scCUT&Tag obtained by scBridge. Colors indicate the Seurat prediction scores and the predicted labels, respectively. B.) Heatmaps representing the overlap score quantifying the similarity upon label transfer between the labels predicted by scBridge and GFP+ G4 scCUT&Tag clusters. The UMAP represents the corresponding Seurat clusters. C.) Heatmap representing the overlap score quantifying the similarity upon label transfer between the scBridge and Seurat predictions. For the labels coming from the Seurat prediction only the cells with prediction score > 0.5 were considered. D.) Representative genome browser track views of potential G4 physical interactions with cicero co-accessibility score > 0.5. E.) Genome browser view of Klk6 promoter in the unsorted and annotated GFP+ sorted sample, respectively. F.) Representative example of the astrocyte specific G4 enrichment in the proximity of gene Gfap. G.) G4 enrichments of the gene body and promoter of Idh1 in OPC and MOL cells. IGV tracks show RPGC normalized G4 signal.

